# Aging and senescence-associated analysis of the aged kidney glomerulus highlights the role of mesangial cells in renal aging

**DOI:** 10.1101/2023.08.10.552883

**Authors:** Ben Korin, Shimrit Avraham, Andrew McKay, Steffen Durinck, Reuben Moncada, Hari Menon, Spyros Darmanis, Yuxin Liang, Zora Modrusan, Joshua D. Webster, Andrey S Shaw

## Abstract

Most causes of chronic kidney disease begin with injury to the glomerulus and involve progressive loss of kidney function. The glomerulus is a capillary bed where blood filtration to produce urine in the kidney occurs. During aging, there is progressive loss of glomeruli and filtration capacity of the kidney because podocytes, the glomerular epithelial cell, are lost with aging and after injury. Podocytes cannot divide and therefore cannot be replaced. Our histological analysis confirmed the presence of glomerulosclerosis, generalized interstitial fibrosis and glomerular hypertrophy in the aged mouse kidney. One barrier to studies of glomeruli is their low frequency in the kidney, less than 1.5% of the cells, and as such, they are often underrepresented in whole kidney analyses. To address this challenge, we used both bulk and single cell RNA sequencing (scRNA-Seq) to characterize purified glomeruli from young and aged mice. Aged glomeruli displayed increased inflammation and expressed a variety of injury and senescence-associated markers, most notably in mesangial cells and macrophages. This increased expression of senescence markers in mesangial cells of aged kidneys suggests a potential cellular target to address age-related renal dysfunction and chronic kidney disease (CKD), which represent a tremendous unmet medical need.

## Introduction

Chronic kidney disease (CKD) is an epidemic affecting more than 10% of the world’s population^1^. One of the biggest risk factors for chronic kidney disease is aging, as a progressive loss of glomerular function is a normal part of aging^2,3^. Central to the function of the kidney is the glomerulus, a capillary bed where blood is filtered to make urine. The glomerulus is made up of three major cells types, the podocyte, the mesangial cell and the endothelial cell. Because the podocyte is an end-stage cell, and unable to divide, the progressive loss of podocytes most likely explains the progressive loss of glomerular function with age^4^. Stressors to the glomerulus like hypertension, diabetes, and immune mediated inflammation contribute to the risk of CKD by accelerating the loss of podocytes.

Whether constitutive podocyte loss is responsible by itself for decline in renal function with age, or whether age-related processes such as inflammation contribute to glomerular decline is not known. Loss of podocytes leads to a compensatory change where the remaining podocytes hypertrophy to try to maintain the filtration surface area^5^. In addition, the glomerular blood pressure is increased to maintain blood filtration by increasing blood flow. The mesangium of the glomerulus, the stromal stalk that holds the capillary loops together, often changes with aging and glomerular injury^6,7^. Greater numbers of mesangial cells, increased expression of extracellular matrix components, recruitment of macrophages, and acquisition of contractile functions are some of the known changes that mesangial cells undergo when homeostasis is compromised^8,9^. Less is known about glomerular endothelial cells, but changes to endothelial fenestrae have been reported^10^. Thus far, we have limited understanding of the gene expression changes that occur in aged glomerular cells, the contribution of cellular senescence to their loss of function, and the properties of the senescent phenotype in glomerular cells.

Recently, efforts like the Tabula Muris Consortium^11^ have generated detailed single-cell studies of multiple organs in mouse that extend across the lifespan of the animal. However, because glomeruli constitute only a small proportion of the total kidney (<1.5%)^12,13^, these databases contain few, if any, glomerular cells. For example, in the Tabula Muris database, only 2 or 3 cells exhibit podocyte markers. To fill this gap, we focused in our study on aging glomeruli, using a technique that uses magnetic beads to purify glomeruli to enrich their representation in our cell preparations^14^.

Here we characterized the aged glomerulus by histology, bulk RNA-Seq and scRNA-Seq. We validated our transcriptomic findings by histology and found increased fibrosis in the aged kidneys, with glomerular loss (glomerulosclerosis), hypertrophy and inflammation. Bulk RNA sequencing (bulk RNA-Seq) demonstrated clear transcriptional changes between aged and young glomeruli that could be attributed to both changes in cell composition as well as cell intrinsic changes associated with aging. ScRNA-Seq analysis allowed us to analyze transcriptional changes in individual cell types with aging, revealing similarities and differences in the transcriptional aging signature of glomerular cells.

Cellular senescence, characterized by a non-proliferative state, cell cycle and growth arrest, and altered cellular function, is one of the hallmarks of aging. Excessive accumulation of senescent cells has been shown to have detrimental effects on tissue function and contribute to age-related diseases, including kidney dysfunction^15^. Work in the kidney and other organs suggests that the accumulation of cell stresses and injuries, coupled with increased demands for cell proliferation, can induce senescence^16^, also promoting an inflammatory state via the secretion of chemokines and inflammatory cytokines^17,18^. Given the fixed cell composition of the glomerulus and the requirement that cells like podocytes survive for the entire lifetime of the animal, we hypothesized that cellular senescence could contribute, along with other CKD risk factors, to glomerular decline. Thus, we utilized our scRNA-Seq dataset to identify and characterize glomerular cells that presented with a senescence-associated transcriptional phenotype. We found that mesangial cells are the cells responsible for the bulk of senescence associated gene expression in the glomerulus. As such, mesangial cells may represent a cellular target of CKD therapeutics.

## Results

### Increased fibrosis, glomerulonephritis, glomerular size, and expression of senescence-associated markers in the aged kidney

First, to confirm and characterize the presence of age-related changes in the kidney of aged mice we performed histology comparing young (3 month-old) and aged (22-24 month-old) mouse kidneys. Kidneys of the aged group were characterized by significantly increased levels of fibrosis (**Figure 1a**) and glomerulonephritis (**Figure 1b**). Extended analysis of 6, 12, 18 and 24 month-old mice identified that both fibrosis and glomerular score progressed with age (**Figure 1c,d**). When glomerular size was measured, both cortical and juxta-medullary aged glomeruli had a larger circumference on average (**Figure 1e**). These results are in-line with previous data of the aged kidney^3,19,20^.

**Figure 1:**
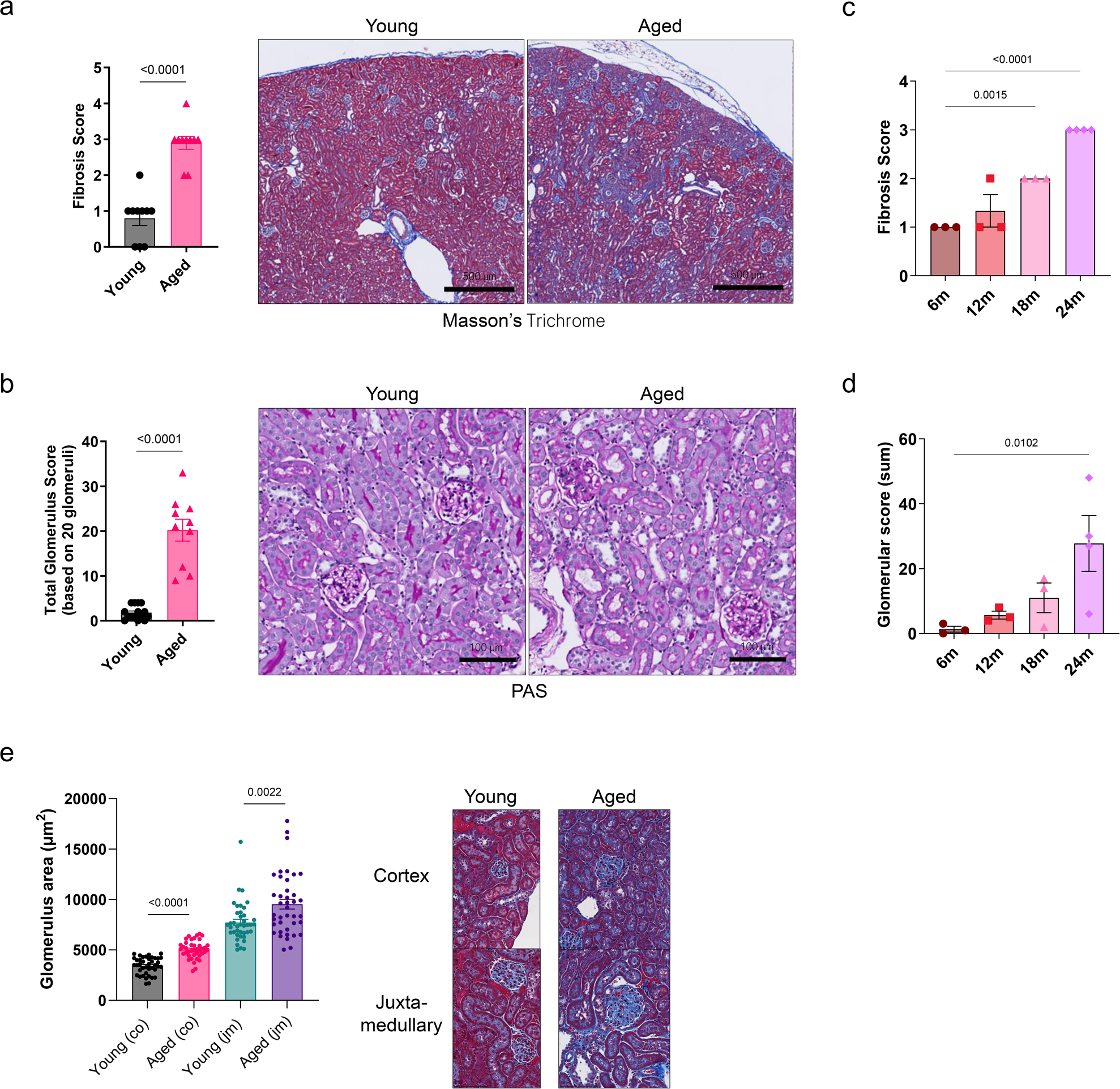
Age-related effects on the kidney and glomerulus. (**a**) Fibrosis (Masson’s Trichrome staining) and (**b**) glomerulonephritis (PAS staining) score analyses of young and aged kidney samples of male mice (n=10,10) with representative images. (**c,d**) Fibrosis and glomerular score by age (n=3,3,3,4 male mice). (**e**) Glomerular size analysis (circumference) of cortical (co) and juxt-medullary (jm) glomeruli of young and aged mice with representative images (n=4,4 male mice, 10 glomeruli per mouse).

### The transcriptional landscape of the aged glomerulus

To identify the molecular mechanisms that accompany glomerular aging, we isolated glomeruli from young and aged mice and performed bulk RNA-Seq (**Figure 2a**). Young and aged glomerular samples markedly differed in their transcriptional phenotype (**Figure 2b**, **Supplementary Table 1a**). Aged samples had increased expression of multiple genes, such as *Apob*, *Apoe*, *Mmp12, Mmp13, Mmp27, Siglece*, *C1qa*, *C1qc*, and decreased expression of *Nell2*, *Gzma*, *Zdbf2* when compared to young samples (**Figure 2c**, **Supplementary Table 1a**). Gene ontology analysis of the differentially expressed genes suggested the involvement of immune system biological processes and the downregulation of developmental processes in aged glomeruli (**Figure 2d, Supplementary Table 1b,c**). These results prompted us to focus on the expression pattern of extracellular matrix components, cytokines and chemokines, macrophage-related and renal disease, injury and repair genes, revealing the relative higher expression levels of many of those genes in the aged glomerulus (**Figure 2e-h**).

**Figure 2:**
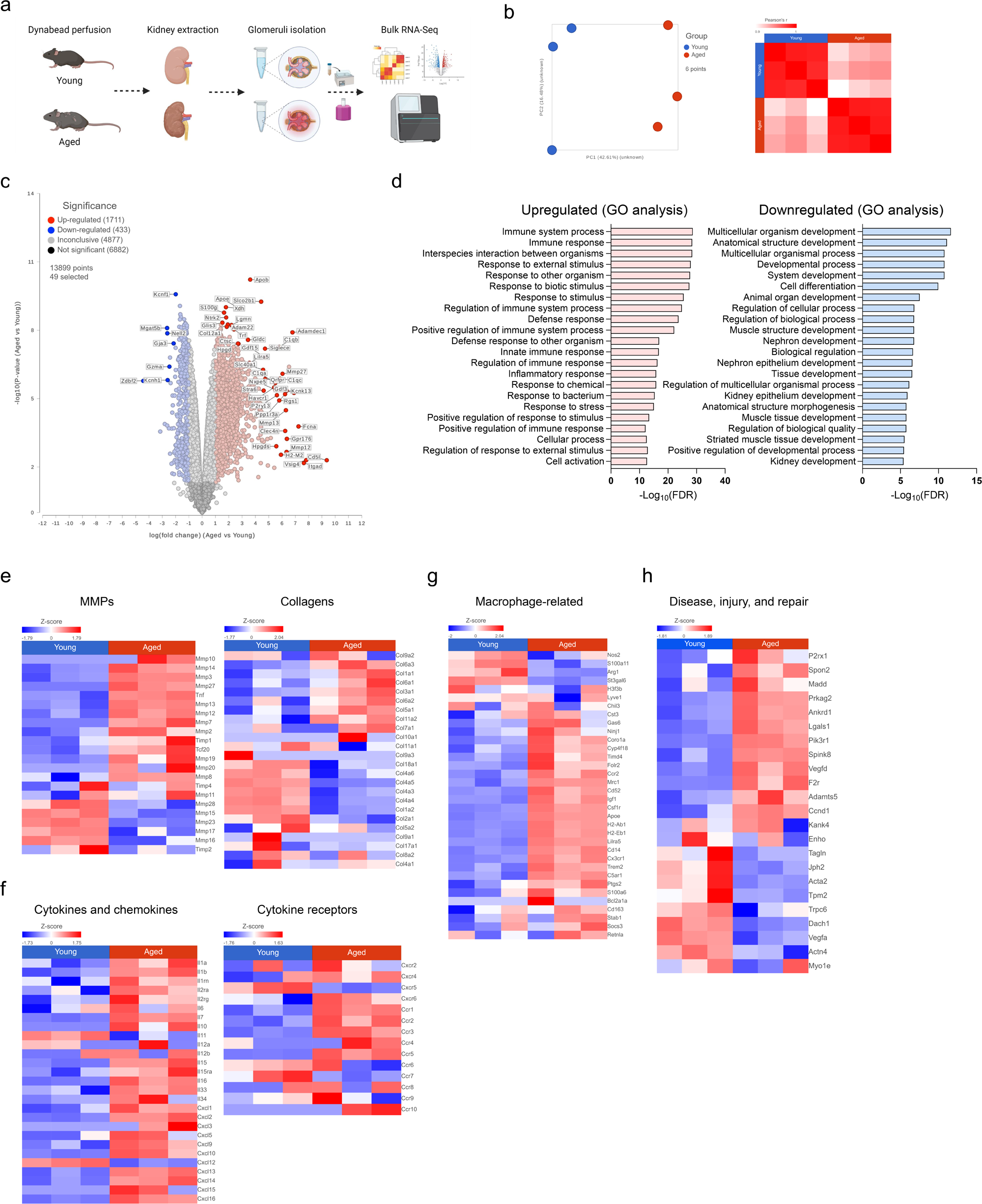
Bulk RNA-Seq analysis of isolated glomeruli reveals induction of pro-inflammatory, ECM, and senescence pathways in the aged glomerulus. (**a**) Illustration of the experimental design for bulk RNA-Seq of young and aged glomeruli. Created with BioRender.com. (**b,c**) PCA plot, correlation matrix, and volcano plot of young and aged glomerular samples according to bulk RNA-Seq data (n=3,3 male mice). (**d**) Biological process strength according to gene ontology (GO) analysis of differentially expressed genes (upregulated or downregulated, genes with > 2 or < (−2) log fold change of expression) in aged glomeruli. (**e-g**) Heatmaps of (**e**) matrix metalloproteases (MMPs) and collagens, (**f**) cytokines, chemokines and their receptors, (**g**) macrophages-related, and disease injury and repair genes according to bulk RNA-Seq data.

### ScRNA-Seq of aged glomerular cells highlights cell type specific age-related changes in endothelial cells, mesangial cells, macrophages, and podocytes

We purified mouse glomeruli and dissociated them into single cell suspensions using a method we previously developed^14,21^, and analyzed them using scRNA-Seq (**Figure 3a**). After unbiased clustering, we assigned cell annotations based on known marker genes. We identified endothelial cells (glomerular capillary and arteriolar), mesangial cells, podocytes, parietal epithelial cells (PECs), vascular smooth muscle cells (SMCs), macrophages, B cells, and T cells (**Figure 3b**, **Supplementary Figure 1a,b**). The separation between young and aged cells in some cell types was evident when colored by group (**Figure 3c**), suggesting that age substantially affects the transcriptional phenotype of certain cell subsets. Young and aged glomerular capillary endothelial cells, arteriolar endothelial cells, mesangial cells, podocytes, and macrophages clustered separately, and differed in their abundance (**Figure 3c**, **Supplementary Figure 1c**). B cells, T cells and SMCs clustered together, suggesting fewer transcriptional differences. Most noticeable was the relative increase in the percentage of macrophages and endothelial cells in aged glomeruli (**Supplementary Figure 1c**). The percentage of vascular SMCs, podocytes, PECs, and B cells was lower with age, while the mesangial cell percentage was similar (**Supplementary Figure 1c**). Macrophages were mostly characterized by high expression of *Cd68, Ninj1, Apoe, H2-Ab1, H2-Eb1 B2m, Cd63*, and *Mif* (**Supplementary Figure 2a**); aged macrophages expressed more *Cd14*, *Socs3*, and *Mrc1* (**Supplementary Figure 2b,c**). To confirm the concordance between our bulk RNA-Seq and our scRNA-Seq experiments we pooled the scRNA-Seq data to generate a pseudo-bulk dataset. This largely recapitulated the bulk RNA-Seq data and validates the accuracy of our scRNA-Seq data (**Figure 3c**).

**Figure 3:**
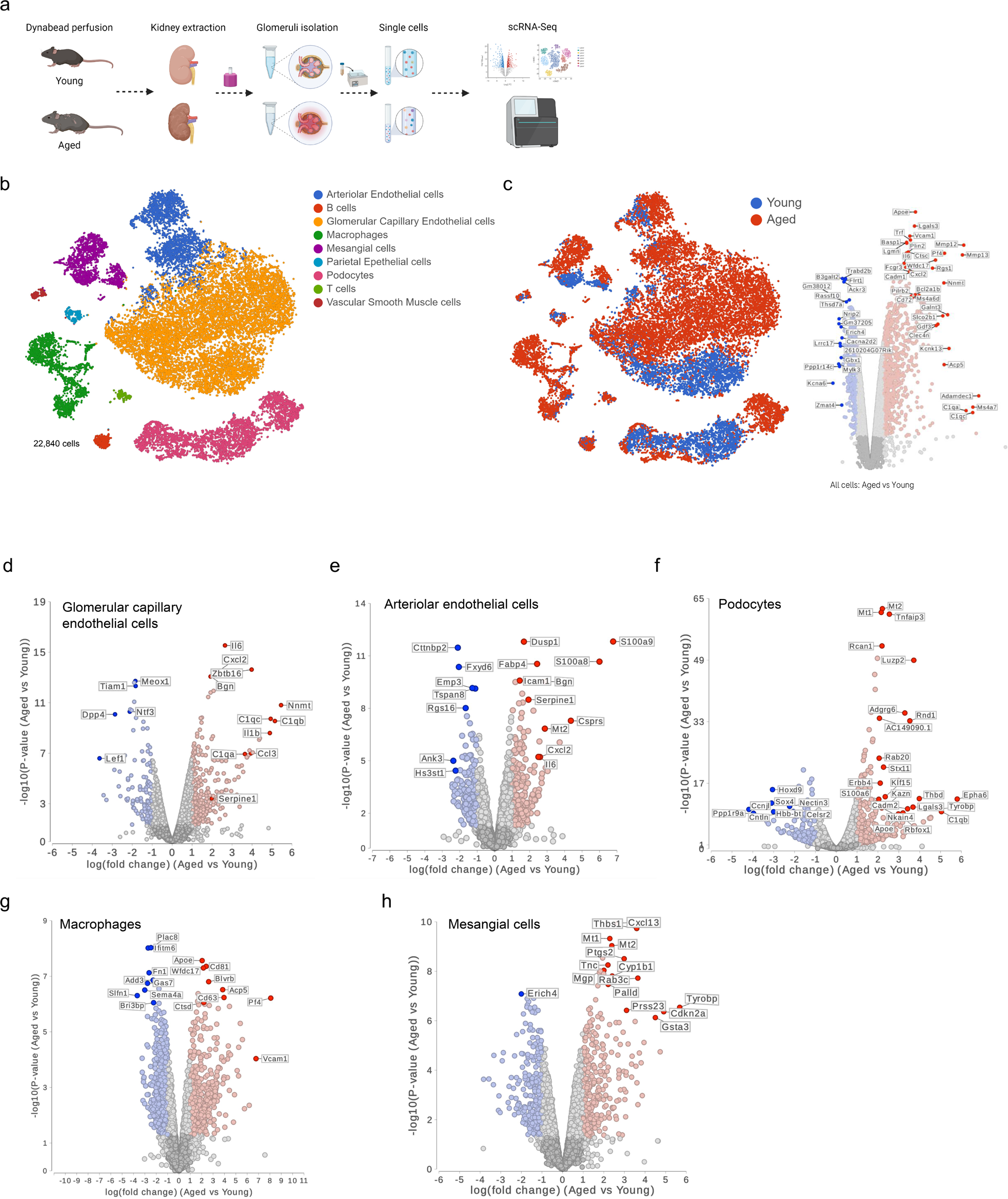
Glomerular scRNA-Seq identifies substantial age-related changes in endothelial cells, podocytes, macrophages, and mesangial cells. (**a**) Illustration of the experimental design for scRNA-Seq of young and aged glomerular cells extracted from male mice. Created with BioRender.com. (**b,c**) tSNE overlay plots by (**b**) identified cell types and (**c**) age group, 22,840 cells in total (7,161 young, 15,679 aged). (**d-h**) Pseudo-bulk cell type specific analysis of aged versus young glomerular cells, volcano plots with highlighted genes in glomerular capillary endothelial cells, arteriolar endothelial cells, podocytes, macrophages, and mesangial cells.

To identify cell type specific age-related transcriptional changes in glomerular cells, we analyzed the differentially expressed genes between aged and young cells by cell type (**Figure 3d-h**, **Supplementary Table 2**). Aged glomerular capillary endothelial cells had increased expression of several cytokines (*Il6*, *Cxcl1*, *Cxcl2*, *Il1b*, *Ccl3*), complement components (*C1qa*, *C1qb*, *C1qc*), and others (*Mt1*, *Mt2*, *Bgn*, *Serpine2*, *Zbtb16*, *Nmt*, *Cd14*, *Lrg1*). Biological process analysis suggested the promotion of chemotaxis, adhesion, remodeling, and inflammatory processes (**Figure 3d**, **Supplementary Figure 3a**). Arteriolar endothelial cells were characterized by increased expression of *S100a8* and *S100a9* genes that play a role in inflammatory process regulation and immune response, and higher levels of the cytokine genes for *Cxcl1*, *Cxcl2*, *Il6*, and others (*Ctla2a*, *Dusp1*, *Icam1*, *Serpine1*, *Smad6*, *Csprs*, *Mt1*, *Mt2*). Biological process analysis suggested the involvement of morphogenesis, collagen biosynthesis, apoptosis, inflammatory, and chemotaxis pathways (**Figure 3e**, **Supplementary Figure 3b**).

Aged podocytes expressed higher levels of metallothioneins (*Mt1, Mt2*)*, Rcan1, Ctgf, Cebbpd, Luzp2, Phlda1, Adgrg6, Apoe, Rab20, Cxcl13*, and complement genes (*C1qb*, *C1qc*) (**Figure 3f)**. Biological process analysis of the significantly upregulated genes suggested the induction of detoxification, adhesion, immune activation, and apoptosis processes (**Supplementary Figure 3c**).

Macrophages had higher expression of *Apoe, Mrc1, Cxcl16, Vcam1, C1qa, C1qc, Cd63, Cd38, Pf4, Igf1, Timp2, Mmp12, Mmp13, Mmp14,* in the aged group (**Figure 3g**). Biological process analysis of the significantly upregulated genes suggested the induction of innate immune response, endocytosis, cytokine, and migration processes (**Supplementary Figure 3d**).

Among other genes, aged mesangial cells had increased expression of the vascular-associated factor *Vcam1,* metallothioneins (*Mt1, Mt2*), cytokines (*Il6, Cxcl13, Ccl2, Cxcl2, Tgfa*), the senescence-associated gene *Cdkn2a* (p16), and extracellular matrix related genes (*Mmp3, Col3a1*) (**Figure 3h**). Biological process analysis suggested the induction of detoxification, cell adhesion and migration, aging, apoptosis, inflammation, cell apoptosis and proliferation pathways (**Supplementary Figure 3e,f**). Additionally, within the various cell types of the aged glomerulus, mesangial cells expressed more genes associated with CKD and glomerular injury and repair^6^ (**Supplementary Figure 4a**). Collagen and MMP gene expression varied between the groups by cell type, noted by an increase in *Col1a2* expression in podocytes, *Mmp12, Mmp13,* and *Timp2* expression in macrophages, and *Col6a3, Col8a1, Col12a1* expression in mesangial cells. Collectively, different cell types of the aged group shared several biological pathways, mostly related to inflammation and immune cell chemotaxis. Yet, this analysis suggested that changes in aged mesangial cells were most related to mechanisms of aging and cell senescence. This encouraged us to further deepen our analysis of the data seeking for senescent cells.

### Analysis of senescence-associated cell signatures suggests mesangial cells as the predominant cell type affected by age and senescence processes

Analysis of the glomeruli bulk RNA-Seq data using known senescence, cell cycle, and SASP genes revealed increased expression of many of the markers in aged glomeruli (**Figure 4a**). Yet, while cell cycle arrest genes were upregulated (e.g., *Cdkn1a*, *Cdkn2a, Cdkn2b*) in the aged group, the expression of the proliferation marker *Mki67* increased in aged as well.

**Figure 4:**
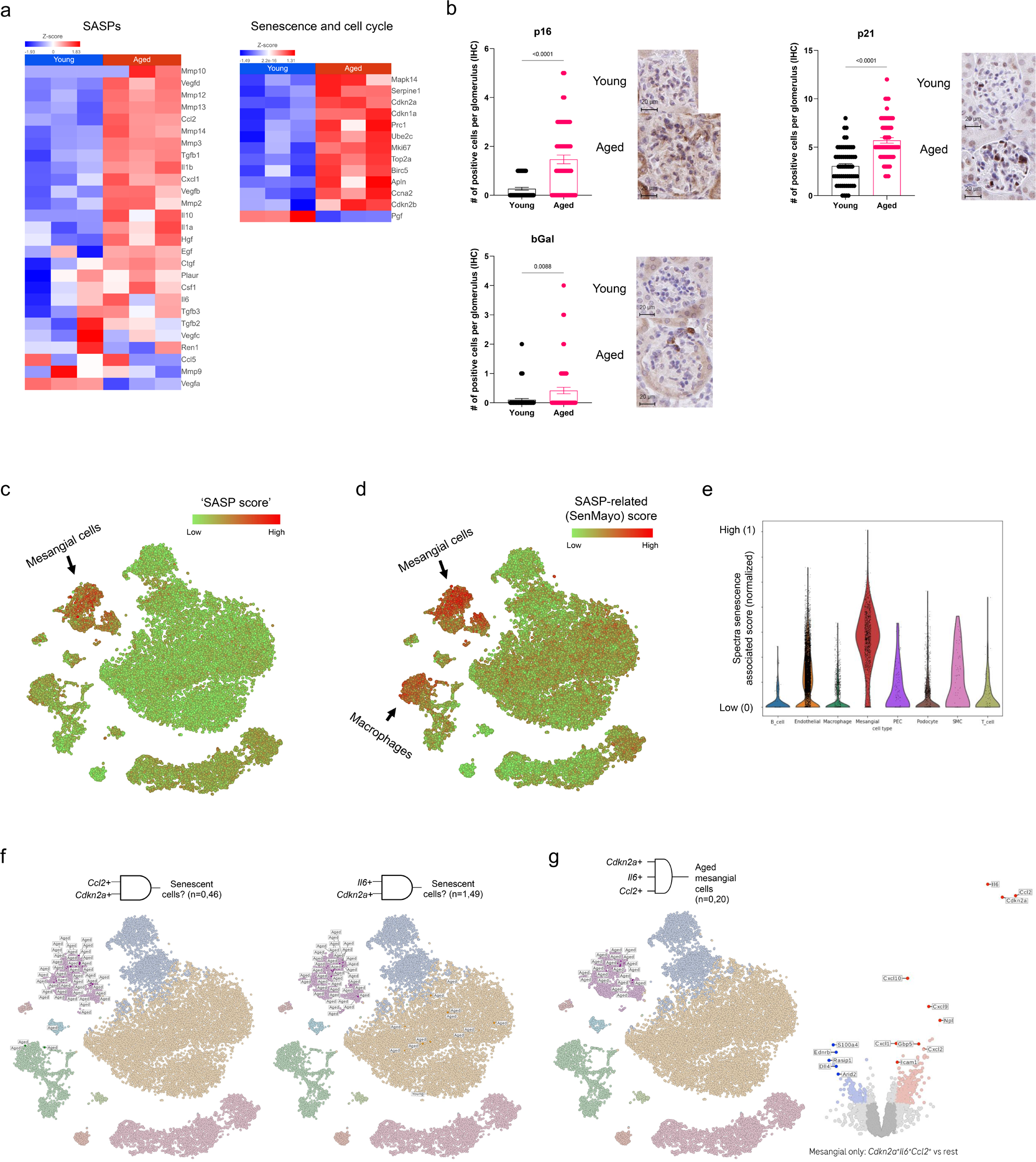
Analysis of senescence-associated markers and signatures highlights mesangial cell senescence in the aged glomerulus. (**a**) Heatmaps of senescence-associated secretory phenotype (SASP) and senescence and cell cycle genes according to bulk RNA-Seq data. (**b**) IHC analysis of p21, p16 and b-Gal staining, the average number of positive cells per was calculated (n=3,3 male mice, 20 glomeruli per mouse). (**c**) ‘SASP score’ and (**d**) SASP-related (SenMayo) score overlay tSNE plots. (**e**) Spectra gene score violin plot. (**f**) Filtered cells expressing both *Ccl2* and *Cdkn2a* (left) or *Il6* and *Cdkn2a* (right) with age annotation. (**g**) Filtered cells expressing both *Cdkn2a*, *Il6*, and *Ccl2*, and volcano plot of differentially expressed genes between *Cdkn2a*^+^*Il6*^+^*Ccl2^+^* mesangial cells and *Cdkn2a^neg^Il6^neg^Ccl2^neg^*mesangial cells.

Staining for known senescence markers: p21, p16, β-Gal (beta-galactosidase), and analyzing the number of positive cells per glomerulus confirmed this phenotype, as aged animals had more glomeruli positive for those senescence markers (**Figure 4b**). However, cells expressing senescence markers by cell staining were relatively rare. The increased expression of cell cycle arrest genes in aged samples of our bulk RNA-Seq analysis suggested the presence of senescent cells in the aged glomerulus. Since the definition of senescent cells *in vivo* is not completely known, we first focused on genes that are associated with a senescence transcriptional signature. Scoring using a previously defined gene set for senescence (SenMayo^22^) was mostly similar between young and aged cells and did not provide significant results (**Supplementary Figure 5a-c**). We therefore first focused on genes associated with secreted factors, using an unbiased approach to interrogate SASP of glomerular cells. Focusing on the senescence-associated secretory phenotype rather than cell cycle arrest, we used a scoring metric of known SASP genes (‘SASP score’). Aged mesangial cells scored highly and had a stronger SASP signature overall compared to other glomerular cell types (**Figure 4c, Supplementary Figure 6a,b, Supplementary Table 3**). As an external impartial metric, an additional SASP-related murine gene list^22^ derived from the SenMayo study was used and showed a higher score in aged mesangial cells and macrophages (**Figure 4d**, **Supplementary Figure 6c,d, Supplementary Table 3**). Using a recently described method for supervised discovery of interpretable gene programs from single-cell data (Spectra^23^), we aimed to refine the SASP-related SenMayo gene signature, and detected additional genes that may be associated with glomerular senescence. Spectra overcomes the cell-type signal dominance by modeling cell-type-specific programs, and this analysis yielded five additional genes (*Marcksl1*, *Bst2*, *H2-K1*, *Selenop*, *B2m*; **Supplementary Table 3**). Scoring for the new signature showed a higher specificity for mesangial cells, which had the highest score of all cell types (**Figure 4e**). The glomerular bulk RNA-Seq analysis confirmed the differential expression of these genes between young and aged glomeruli (**Supplementary Figure 7**).

Next, we focused on genes associated with senescence individually. We identified that some markers were expressed in a cell specific fashion (e.g., *Cdkn1c* (p57)) or were commonly expressed (e.g., *Cdkn1a* (p21)) in both young and aged cells (**Supplementary Figure 8**). Based on their increased expression in aged versus young mice, and their relatively infrequent detection we focused on the cell cycle arrest marker *Cdkn2a* (p16) and the SASP genes *Ccl2* (CCL2) and *Il6* (IL-6) (**Supplementary Figure 8**). We first screened for cells that co-express *Cdkn2a* and *Ccl2* or *Il6*, and found 46 and 49 cells, respectively, that were in the aged group (**Figure 4f**). When we searched for co-expression of all three genes, *Cdkn2a*, *Ccl2*, and *Il6*, 20 cells from the aged animals, all mesangial cells, showed combined expression of all three markers (**Figure 4g**). Differential expression analysis of these 20 cells compared to the remainder of the mesangial cells (negative for *Cdkn2a*, *Ccl2*, and *Il6*) identified other upregulated genes in these cells and potentially represent their senescence signature (**Figure 4g**, **Supplementary Table 3**). Among these genes were *Cxcl1, Cxcl2, Cxcl9, Cxcl10,* that encode for secreted cytokines that may be specifically involved in mesangial cell senescence and aging processes. This approach highlighted that only mesangial cells expressed a combination of these genes. Lastly, the identification of the cellular senescence phenotype of aged mesangial cells was further supported by KEGG cellular senescence pathway analysis that presented an induction in several parameters of this pathway (**Supplementary Figure 9a,b**).

## Discussion

Aging of the kidney is comprised of structural and functional changes with inflammation and fibrosis as the major features^24^. Aging is also associated with a decline in the regenerative capacity of renal cells, compromising the kidney’s ability to repair and regenerate damaged tissue^25^. Oxidative stress, chronic inflammation, genetic and epigenetic factors, telomere dysfunction, and comorbidities such as diabetes and hypertension can accelerate cellular senescence in the kidney^26–29^. This reduced regenerative potential and increase cellular senescence can contribute to the progression of renal disease.

Studies have implicated the connection between increased senescence within the kidney and the progression of CKD^15^. Since injury and loss of glomeruli are central to the development of CKD, we focused on characterizing transcriptional changes associated with aging in purified glomeruli. Because of their low abundance, previous studies of aged kidneys are dominated by changes in tubules as they represent the most abundant cells in the kidney and are lacking in glomerular cells. In our studies, we compared 3-4 month-old animals to 22-24 month-old animals. The aged animals exhibited glomerulosclerosis, glomerular hypertrophy, interstitial fibrosis, and associated inflammation. While comparisons to human are inexact, 22-24 month-old mice are thought to be analogous to 56-69 year old humans.

As glomeruli represent less than 1.5% of the cells in the kidney, our strategy was to first purify glomeruli from kidneys using a protocol based on the use of magnetic beads^14^. Any purification strategy has the potential for the induction of disassociation-associated genes. To minimize such effects, we combined bulk RNA-Seq with scRNA-Seq. While both require tissue disassociation, bulk RNA-seq does not require single-cell disassociation, thus significantly shortening the protocol. Having both bulk and scRNA-Seq data allowed us to directly compare them by aggregating the scRNA-seq data to generate a pseudo-bulk RNA-Seq dataset. We found that the two datasets were comparable. Our protocol was previously optimized for cell viability (>90%) and low mitochondrial counts (<5%). This is an advantage from whole kidney studies, where mitochondrial RNA contamination can be as high as 50%.

As cellular senescence is thought to contribute to diseases of aging, we were very interested to identify markers of senescence in glomerular cells of aging mice. The accumulation of senescent cells in aged glomeruli could potentially be a significant contributor to the development of CKD in humans. However, since most of what is known about senescence is from *in vitro* studies of cultured cell lines, cell senescence signatures *in vivo* in primary cells is an open question.

Encouragingly, many of the genes that we identified as differentially expressed between young and old glomeruli were known senescence genes. Therefore, when we analyzed our scRNA-Seq data we first focused on determining which cells expressed senescence genes that were detected in the bulk RNA-Seq differential gene analysis comparing young and old glomeruli. This suggested that most of the expression was derived from mesangial cells and macrophages. As macrophages in the kidney are generally replenished with bone marrow derived monocytes^30^ and the senescence genes expressed were cytokines and chemokines that they normally express, we focused on mesangial cells.

Analysis of mesangial cells showed that the differentially expressed senescence genes were only expressed in a minority of the cells and the signal was distributed across the population with little overlap. We therefore focused on finding cells expressing two or three of these genes. We found that there were variable numbers of aged mesangial cells that co-expressed *Cdkn2a* and/or *Il6* and *Ccl2*. Importantly, these three genes are canonical genes for cell senescence. Further analysis showed that these cells were distinct for the expression of *Cxcl13* which has been suggested to be involved in the pathogenesis of systemic lupus erythematosus^31^. Given the low transcriptional sampling obtained by scRNA-seq, the number of cells that we identified is almost certainly an underestimate. Purification or enrichment strategies for mesangial cells from young and aged samples could be a way to obtain further information regarding the senescent properties of these cells.

Using a scoring system based on senescent markers and SASPs, we were unable to identify potential senescent cells in any other cell type. There are several possible explanations. First, it is possible that the 22-24 month-old mice were not old enough or required some injury to accelerate the changes of senescence. Second, it is possible that the senescence signature of glomerular cells is unrelated to that defined for other tissues. For example, as podocytes are already cell cycle arrested, the senescence signature could be unique. We did identify that aged podocytes and aged endothelial cells had a higher expression of *C1qa*, *C1qb*, or *C1qc* suggesting a role for complement activation in aging. Third, the number of senescent cells could be very rare and our methods are not sensitive enough to identify them. Alternatively, it is also possible that these cell lineages do not develop a senescence phenotype. It is interesting to note that some of the genes associated with senescence are also genes associated with cell stress and cell disassociation^6,32^.

Previous studies showed that in glomerular endothelial cells, plasminogen activator inhibitor-1 (PAI-1), encoded by the *Serpine1* gene and a known senescence associated gene, can drive age-related kidney disease^33^. Our data confirmed an increase in *Serpine1* expression in aged glomerular capillary and arteriolar endothelial cells, along with other senescence-associated genes (e.g., *Il6*, *Il1b*, *Cxcl2*) but few if any co-expressing cells were identified. Overall, these changes in the various compartments and cell types of the kidney contribute to renal inflammation, fibrosis, and impaired regeneration, promoting the age related decline in kidney function. Addressing cell senescence and tissue regeneration in the kidney holds promise for therapeutic interventions by targeting senescent cells through, for example, senolytic drugs, that selectively eliminate senescent cells^34^. Studies have demonstrated the beneficial effects of senolytic drugs in improving renal function and attenuating fibrosis^35^. Another approach has been focusing on cell reprogramming, using the reprogramming factors Oct4, Sox2, Klf4 and c-Myc for short or long intervals of time, to promote a youthful epigenetic signature, which has showed encouraging results^36^.

Together, our results suggest that age mostly affects mesangial cells and promotes a senescence-associated phenotype in these cells. As a key component of glomerular function, it is plausible that these changes in mesangial cells drive the age-related phenotype within the aged glomerulus, further promoting kidney dysfunction with age. Future studies may benefit from targeting mesangial cells specifically to attenuate the consequences of renal aging.

## Methods

### Animals

C57BL/6J male mice (Jax: 000664), 3-4 months of age (Young) or 22-24 months of age (Aged), weighing at least 20 g or 30-35 g respectively, were used. All animals were housed at Genentech under specific pathogen-free conditions with free access to chow and water and 12/12 hr light/dark cycle. Sample sizes were chosen based on similar experiments that have been previously published. All animal procedures were conducted under protocols approved by the Institutional Animal Care and Use Committee at Genentech, and were performed in accordance with the Guide for the Care and Use of Laboratory Animals.

### Mouse kidney glomeruli isolation and dissociation to single cell suspensions

Mouse glomeruli isolation and preparation of single cell suspensions was performed as shown previously with slight modifications^14^. Briefly, mice were perfused transcardially with 10 ml of RPMI with 200 ul of M-450 Dynabeads (Thermo Fisher Scientific, 14013). Kidneys were immediately extracted and minced using a razor blade, then transferred to 5 ml digestion buffer with Liberase^TM^ (1.5 U/ml; 5401127001, Millipore Sigma) and DNaseI (100 U/ml; D4527, Millipore Sigma) in RPMI-1640, and incubated for enzymatic digestion (37⁰C, 800 rpm, 20 min). Following digestion, samples were sieved via a 100 μm and a 70 μm MACS^®^ SmartStrainers (130-098-463, 130-098-462, Miltenyi Biotec), and 30 ml ice-cold 10% FBS in RPMI-1640 were added to stop digestion. Samples were washed twice with 30 ml ice-cold RPMI-1640 and glomeruli were isolated using a magnetic separator (DynaMag-2, Thermo Fisher Scientific, 12321D). Glomeruli were dissociated in 1 ml digestion buffer with Trypsin (0.5% wt/vol; Thermo Fisher Scientific, 15090046), Dispase II (2 U/ml; Roche, 04942078001), Collagenase D (2 U/ml; Roche, 11088866001), and DNAse I (10 U/ml; D4527, Millipore Sigma), in RPMI-1640, and incubated for enzymatic digestion (37⁰C, 800 rpm, 40-60 min). Samples were kept on ice until further processing. Cell counts and viability were determined before further analyses and viability was determined at >95% viable cells.

### Bulk RNA sequencing (bulk RNA-Seq) and analysis

For bulk RNA sequencing, glomeruli were placed directly into RNeasy Lysis Buffer (Qiagen). RNA was extracted and isolated from samples using RNeasy Mini Kit (74104, Qiagen) according to manufacturer instructions. Regular input RNA-Seq was performed as follows. Total RNA was quantified with Qubit RNA HS Assay Kit (Thermo Fisher Scientific) and quality was assessed using RNA ScreenTape on 4200 TapeStation (Agilent Technologies). For sequencing library generation, the Truseq Stranded mRNA kit (Illumina) was used with an input of 100 nanograms of total RNA. Libraries were quantified with Qubit dsDNA HS Assay Kit (Thermo Fisher Scientific) and the average library size was determined using D1000 ScreenTape on 4200 TapeStation (Agilent Technologies). Libraries were pooled and sequenced on NovaSeq 6000 (Illumina) to generate 30 millions single-end 50-base pair reads for each sample. Data were analyzed and visualized using *Partek^®^ Flow^®^* (https://www.partek.com/partek-flow/).

### Single cell RNA sequencing (scRNA-Seq) and analysis

Samples were kept on ice and immediately processed. Sample processing for scRNA-seq was done using Chromium Next GEM Single Cell 3’ Kit v3.1 (1000269, 10X Genomics) according to manufacturer’s instructions. Cell concentration was used to calculate the volume of single-cell suspension needed in the reverse transcription master mix, aiming to achieve approximately 8000 cells per sample. cDNA and libraries were prepared according to manufacturer’s instructions (10X Genomics). Sequencing reads were processed through 10X Genomics Cell Ranger (v.3.1.0) and aligned to GRCm38. Cells with fewer than 500 genes expressed or more than 5,000 genes expressed and more than 12% mitochondrial content were filtered out. Features (genes) where value <= 0.0 in at least 99.5% of the cells were excluded. Initial cell count: 33,323 cells, after QA/QC: 22,840 cells were analyzed (7,161 young, 15,679 aged). Data were analyzed and visualized using *Partek^®^ Flow^®^* (https://www.partek.com/partek-flow/).

### Histology and pathology analysis

Tissues used for histologic analysis and immunohistochemistry (IHC) were routinely fixed in formalin, paraffin embedded, and sectioned at approximately 4-5 µm. Mouse tissues were histologically scored in a random, blinded manner on hematoxylin and eosin stained sections. The scoring system included assessments of inflammation (0: no significant inflammatory infiltrates; 1: 1-2 small leukocyte aggregates (less than approximately 10 cells or scattered leukocytes associated with adjacent tubular atrophy); 2: 3-5 small leukocyte aggregates or 1 larger aggregate; 3: 6-10 small leukocyte aggregates or 2-3 larger aggregates; 4: >10 small leukocyte aggregates or >3 larger aggregates) and subcapsular tubular atrophy and fibrosis (0: no subcapsular tubular atrophy; 1: 1-2 foci of subcapsular tubular atrophy; 2: 3-5 foci of subcapsular tubular atrophy; 3: 6-10 foci of subcapsular tubular atrophy; 4: >10 foci of subcapsular tubular atrophy). These scores were summed to generate a final score.

### Statistical analyses and reproducibility

Data are presented as mean ± s.e.m. Experiments were repeated independently at least two times (biological replicates). Statistical significance was determined using two-tailed unpaired t-test (nonparametric) when comparing two independent groups. Statistical analyses were performed using Prism 9.5 (GraphPad software). *P values* are provided within figures.

### Data availability

The data that support the findings of this study are available upon reasonable request. The sequencing data generated for this study are available in the Gene Expression Omnibus under accession number <u>GSE240375</u>.

## Acknowledgements

We thank and our Genentech colleagues in the Research Biology, Laboratory Animal Resources, and Pathology Departments for their support in this study. We thank Dr. Mary Mohrin for helpful discussions and insightful comments. This work was supported by Genentech.

## Contributions

B.K. designed and performed experiments, analyzed and interpreted the data, created the figures and wrote the manuscript. S.A. took part in optimization of glomeruli isolation and dissociation method, performed experiments, assisted in data interpretation and revised the manuscript. A.M. supported scRNA-seq data analysis, performed Spectra analysis, and revised the manuscript. R.M., H.M, S.D. Y.L. Z.M. supported bulk RNA-Seq or scRNA-Seq. J.D.W. performed pathology analysis and data interpretation, and revised the manuscript. A.S. oversaw data analysis and interpretation, and revised the manuscript.

## Supplementary data legend

**Supplementary Figure 1: Supporting scRNA-Seq data.** (**a**) tSNE plot by cluster. (**b**) tSNE plots with marker expression overlay. (**c**) Percentage of cells per group, scRNA-Seq data.

**Supplementary Figure 2: Macrophage markers, scRNA-Seq data.** (**a**) tSNE plots of macrophage-related markers. (**b**) Bubble plots of young and aged macrophage markers, scRNA-Seq data.

**Supplementary Figure 3: Upregulated biological processes (gene ontology) in aged glomerular cell types.**

**Supplementary Figure 4: Bubble plots (injury and repair, collagens, MMPs) of young and aged per cell type.**

**Supplementary Figure 5: Supporting data of SenMayo score.** (**a**) Overlaid tSNE plot of SenMayo score. (**b**) Bubble plots of SenMayo score genes in young and aged glomerular cells, scRNA-Seq data. (**c**) Heat map of SenMayo genes in young and aged glomeruli, bulk RNA-Seq data.

**Supplementary Figure 6: Senescence analysis score plots.** (**a,b**) SASP and (**c,d**) SASP-related (SenMayo) bubble plots (scRNA-Seq) and heatmaps (bulk RN-Seq).

**Supplementary Figure 7: Spectra gene expression in glomeruli, bulk RNA-Seq.** Heat map of differential expression of the identified Spectra genes in isolated young and aged glomeruli, bulk RNA-Seq.

**Supplementary Figure 8: Gene expression tSNE plots of senescence-associated genes.**

**Supplementary Figure 9: Cellular senescence KEGG pathways in mesangial cells.** KEGG pathway analysis in (**a**) aged versus young or (**b**) senescent versus non-senescent mesangial cells

**Supplementary Table 1: Differentially expressed genes of aged versus young glomeruli, bulk RNA-Seq.**

**Supplementary Table 2: Differentially expressed genes of aged versus young glomerular cells, per cell type, scRNA-Seq.**

**Supplementary Table 3: SenMayo, SASP, and Spectra gene lists.**

