## Supplementary Figures for "Aging and senescence-associated analysis of the aged kidney glomerulus highlights the role of mesangial cells in renal aging"

Supplementary Figure 1: Supporting scRNA-Seq data.

a

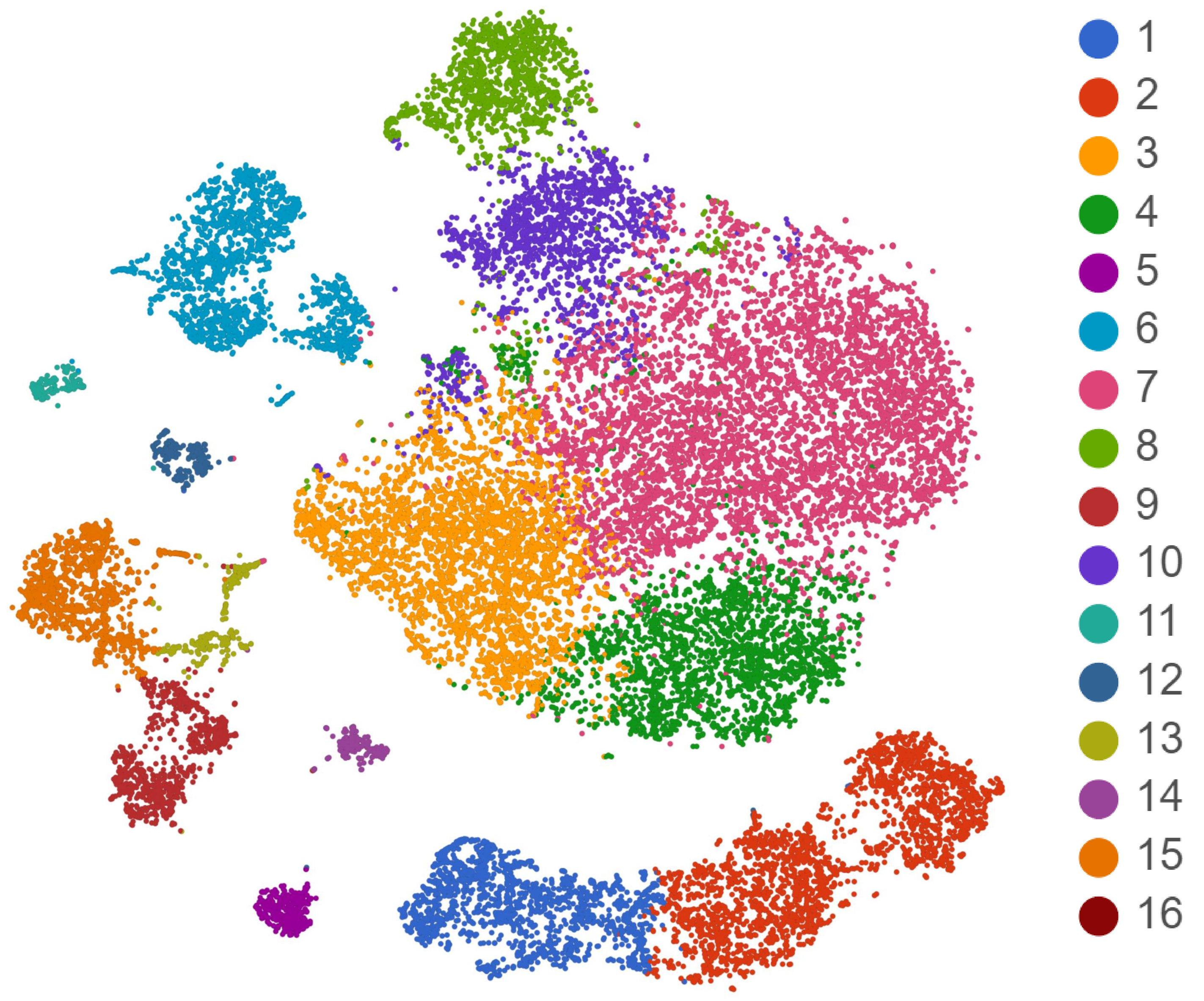


b

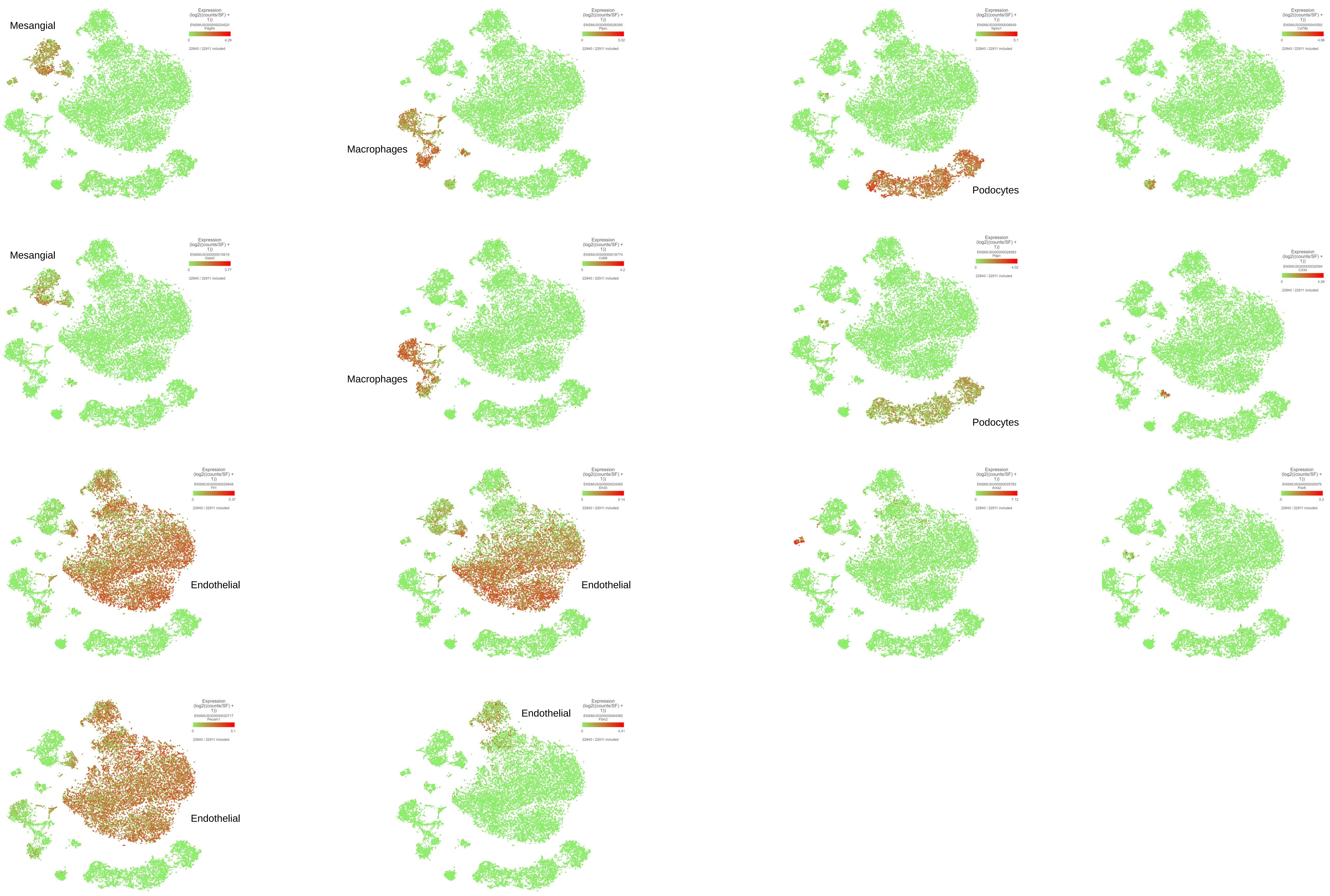


c

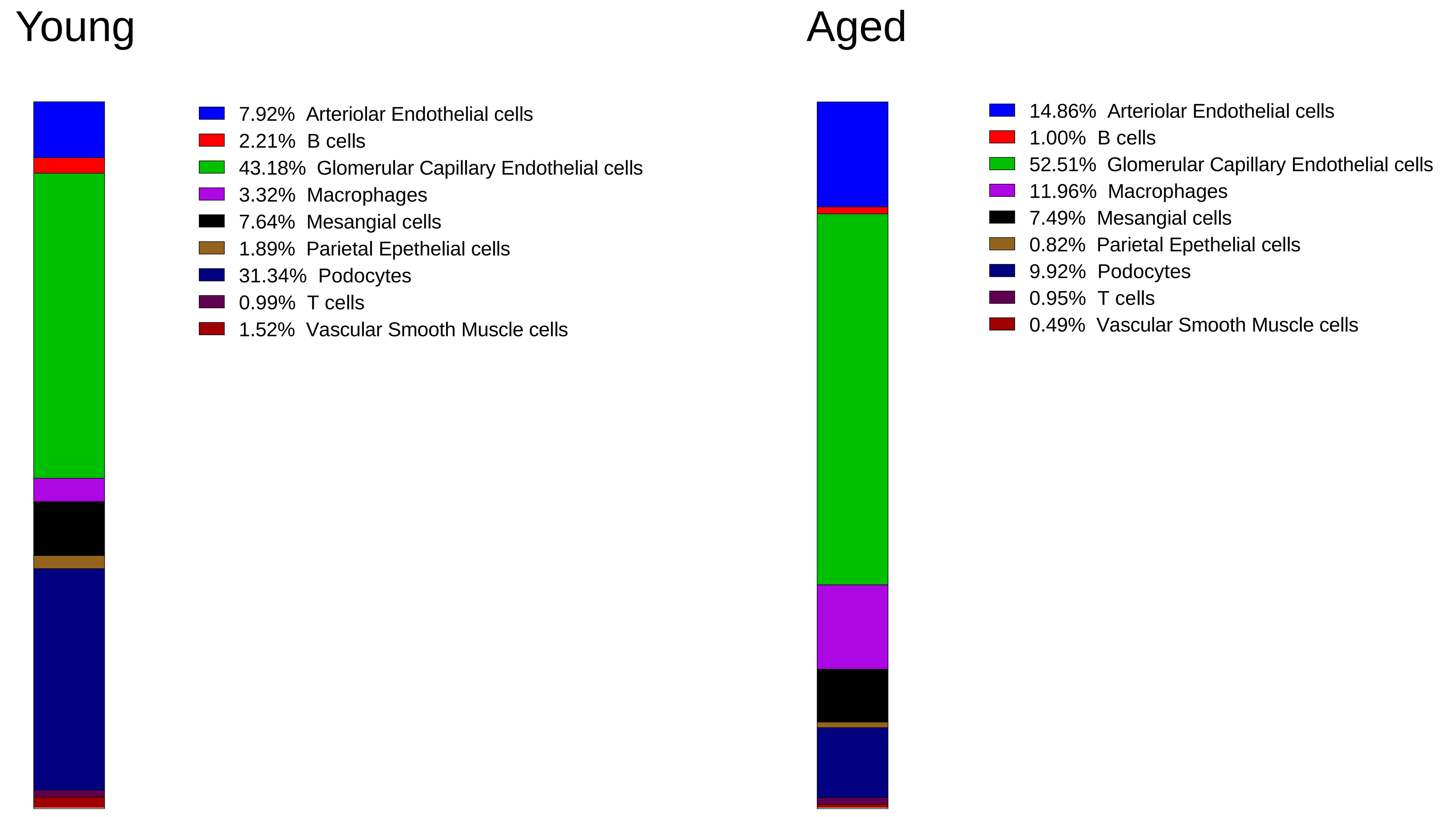

Supplementary Figure 2: Macrophage markers, scRNA-Seq data.

a

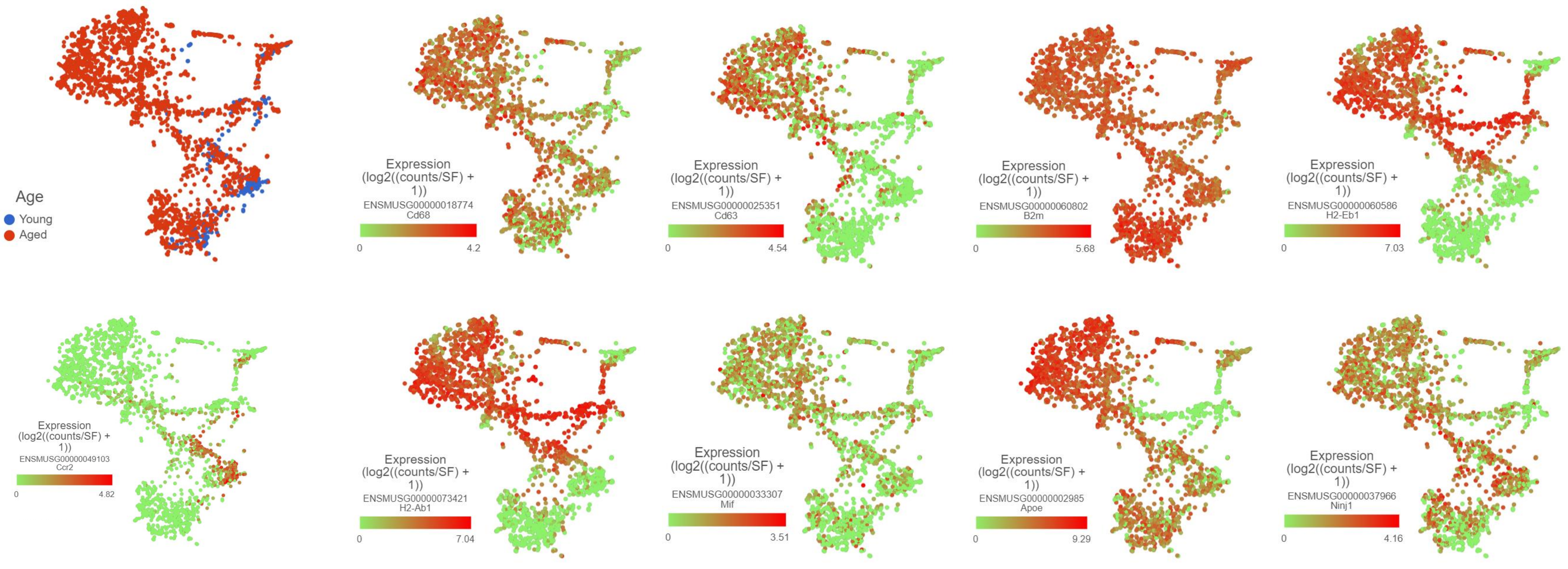


b

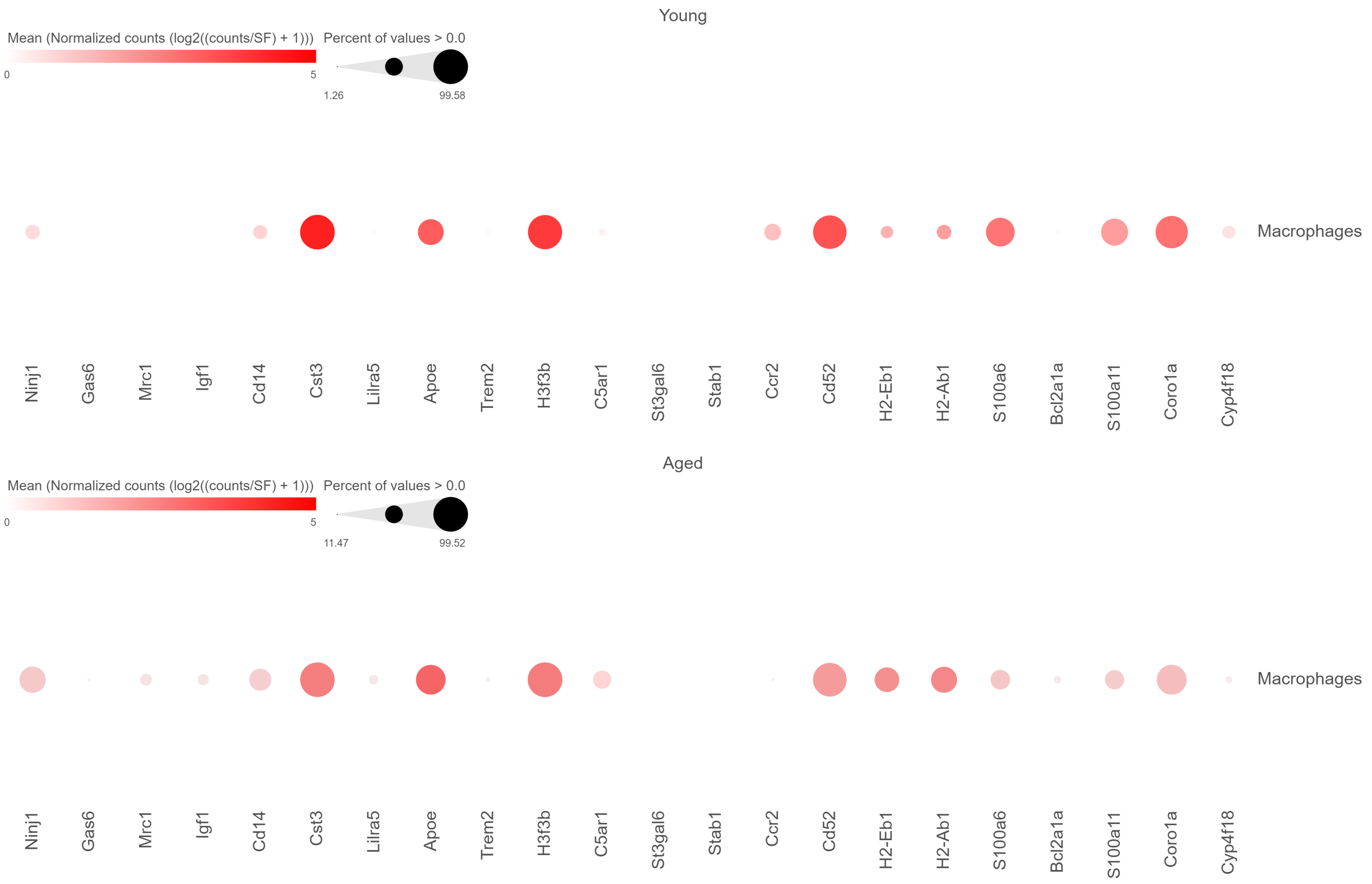

Supplementary Figure 3: upregulated biological processes (gene ontology) in aged glomerular cell types.

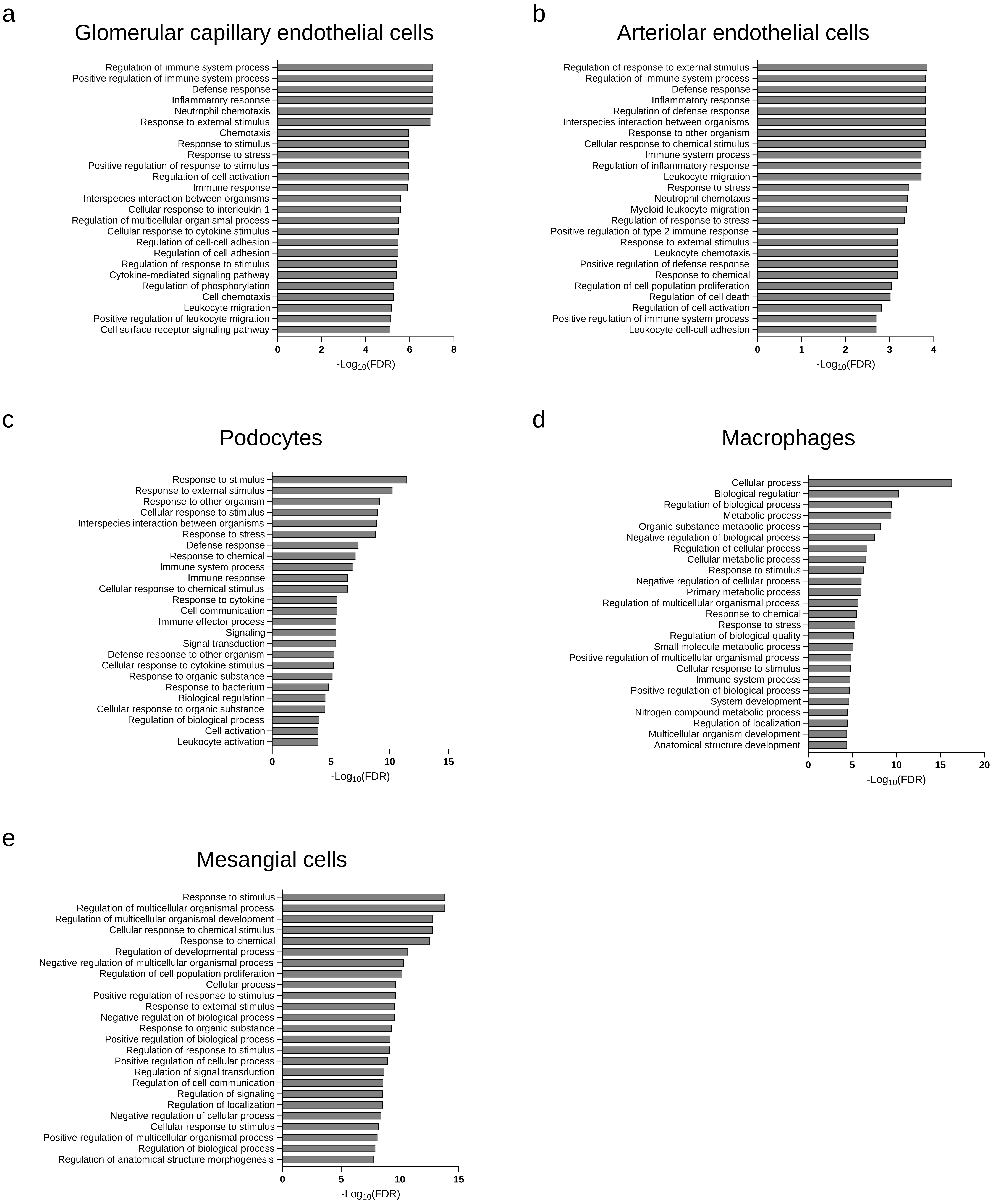

Supplementary Figure 4: Bubble plots (injury and repair, collagens, MMPs) of young and aged per cell type.

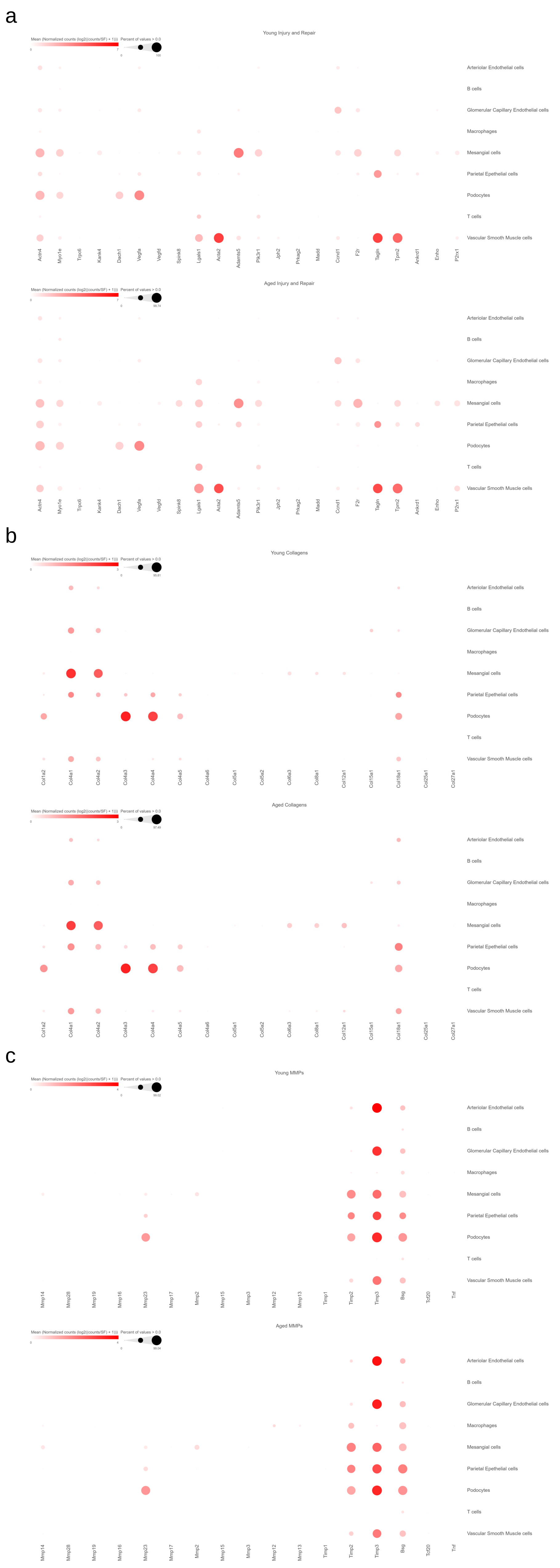

Supplementary Figure 5: Supporting data of SenMayo score.

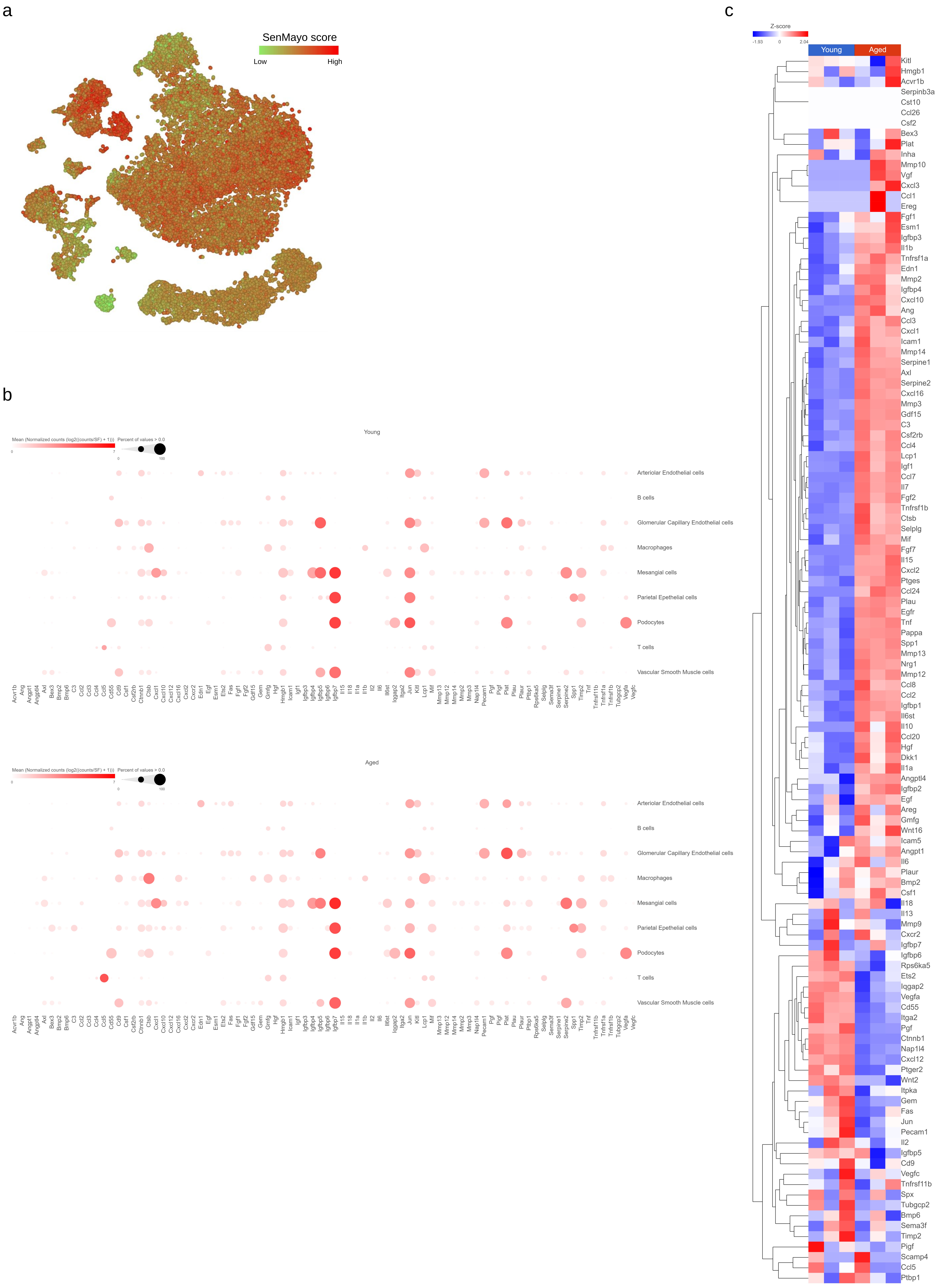

Supplementary Figure 6: Senescence analysis score plots.

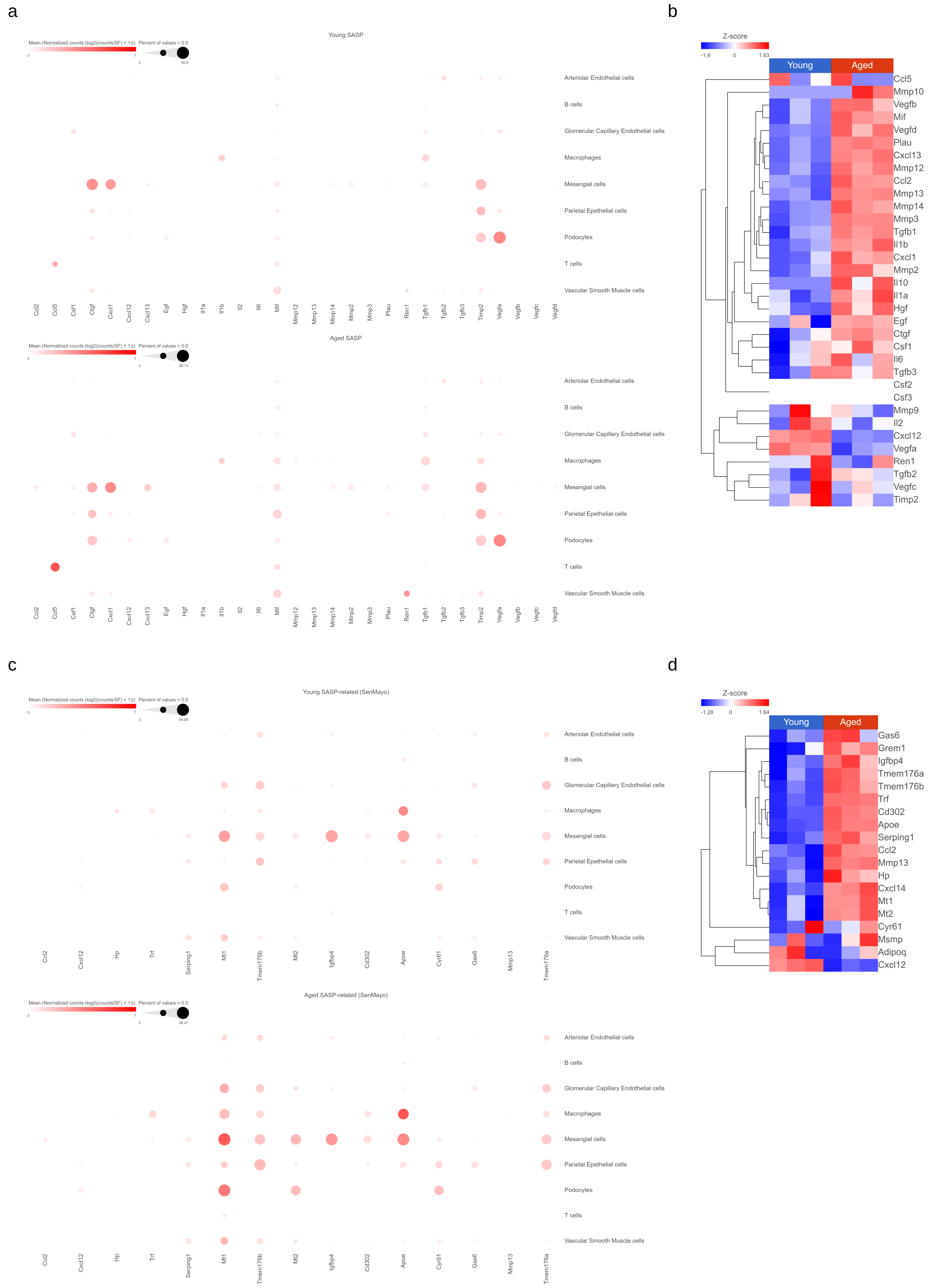

Supplementary Figure 7: Spectra gene expression in glomeruli, bulk RNA-Seq.

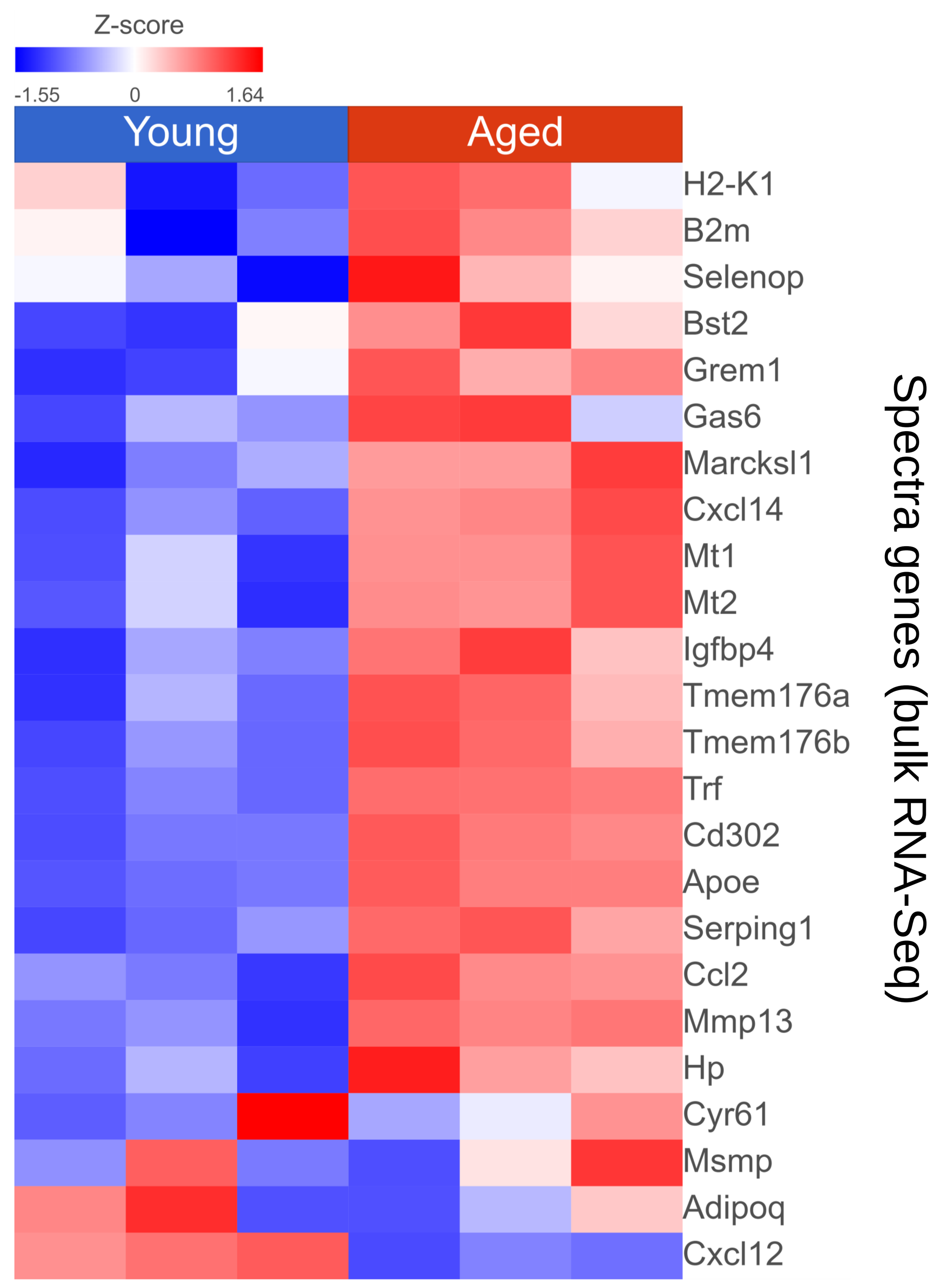

Supplementary Figure 8: Gene expression tSNE plots of senescence-associated genes.

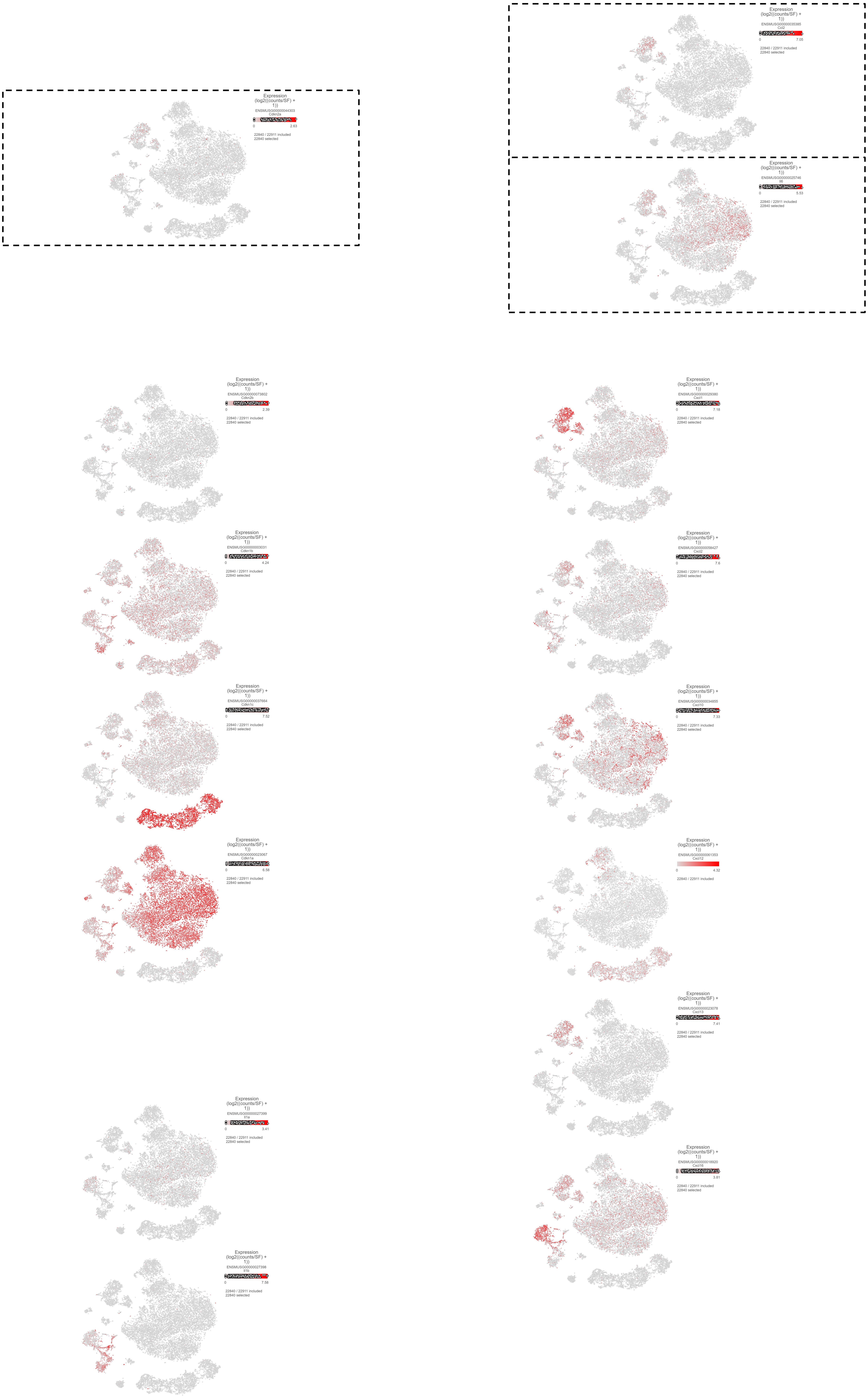

Supplementary Figure 9: Cellular senescence KEGG pathways in mesangial cells.

a

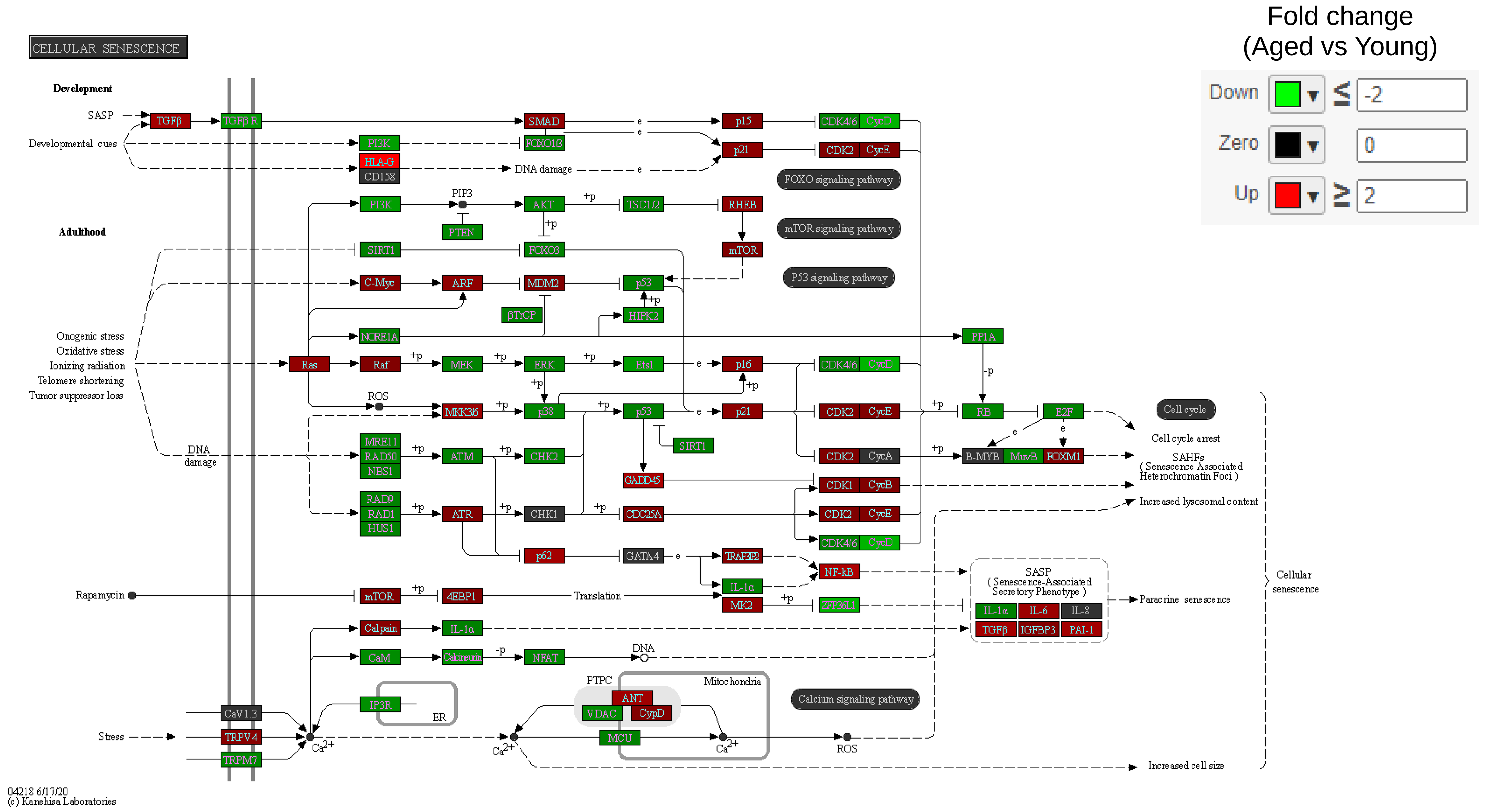


b

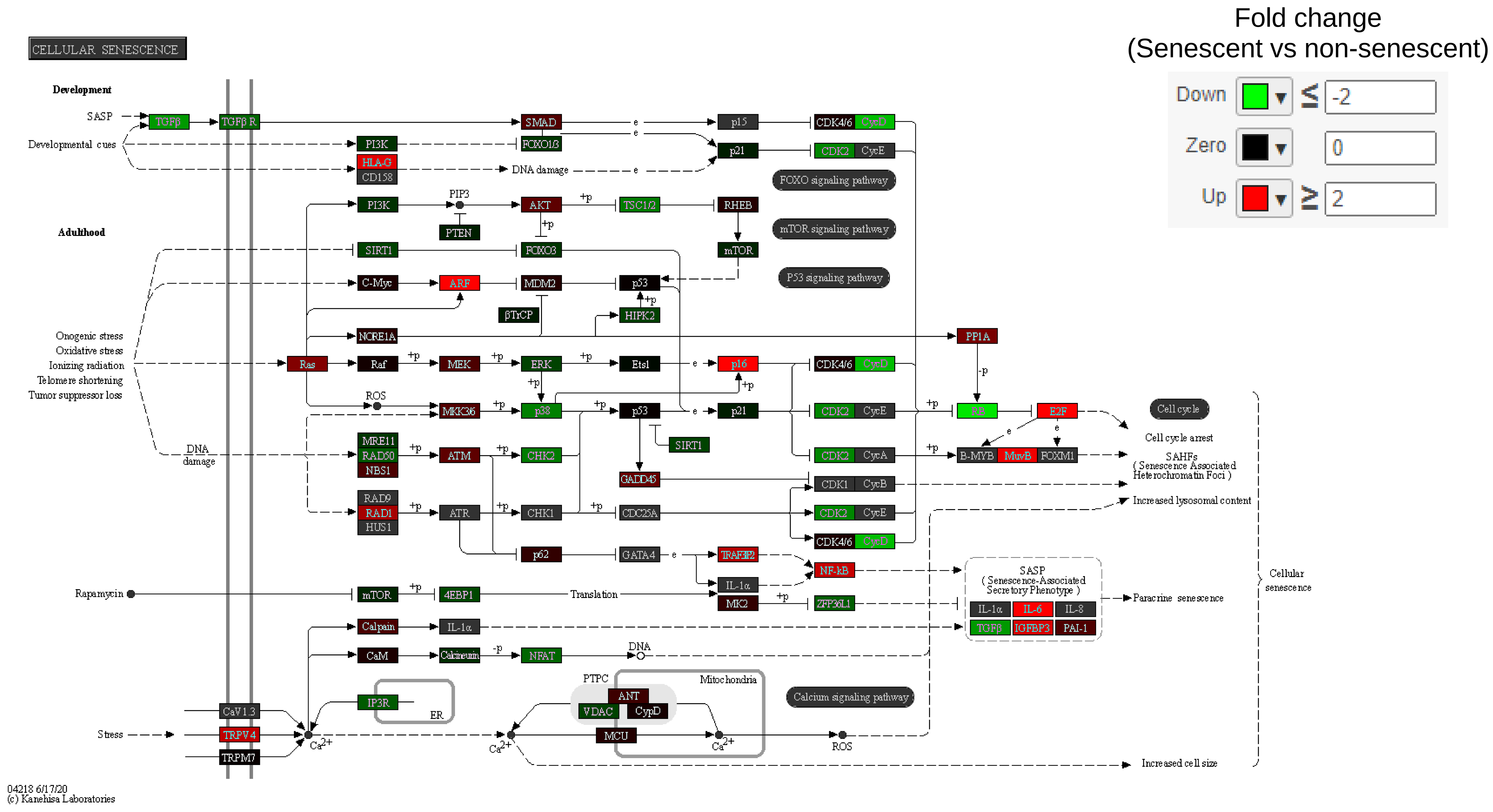
